# Starvation resistance in the nematode *Pristionchus pacificus* requires a conserved supplementary nuclear receptor

**DOI:** 10.1101/2023.08.21.554071

**Authors:** Tobias Theska, Tess Renahan, Ralf J. Sommer

**Author notes:** Corresponding author Ralf J. Sommer.

## Abstract

Nuclear hormone receptors (NHRs) are a deeply-conserved superfamily of metazoan transcription factors, which fine-tune the expression of their regulatory target genes in response to a plethora of sensory inputs. In nematodes, NHRs underwent an explosive expansion and many species have hundreds of *nhr* genes, most of which remain functionally uncharacterized. However, recent studies elucidated that two sister receptors, *Ppa-*NHR-1 and *Ppa-*NHR-40, are crucial regulators of feeding-structure morphogenesis in the diplogastrid model nematode *Pristionchus pacificus*. In this study, we functionally characterize *Ppa-*NHR-10, the sister paralog of *Ppa-*NHR-1 and *Ppa-*NHR-40, aiming to reveal whether it too regulates aspects of feeding-structure development. We used CRISPR/CAS9-mediated mutagenesis to create knock-out mutations of this receptor and applied a combination of geometric morphometrics and unsupervised clustering to characterize potential mutant phenotypes. However, we found that *Ppa-*NHR-10 does not affect feeding-structures morphogenesis. Instead, multiple RNA-seq experiments revealed that many of the target genes of this receptor are involved in lipid catabolic processes. We hypothesized that their mis-regulation could affect the survival of mutant worms during starvation, where lipid catabolism is often essential. Indeed, using novel survival assays, we found that mutant worms show drastically decreased starvation resistance, both as young adults and as dauer larvae. We also characterized genome-wide changes to the transcriptional landscape in *P. pacificus* when exposed to 24hrs of acute starvation, and found that *Ppa-*NHR-10 partially regulates some of these responses. Taken together, we were able to demonstrate that *Ppa-*NHR-10 is broadly required for starvation resistance and regulates different biological processes than its closest paralogs *Ppa-*NHR-1 and *Ppa-*NHR-40.

## Background

Phenotypic plasticity is the ability of a single genotype to produce different phenotypes depending on the environmental cues that the organism receives during development. As such, phenotypic plasticity has re-gained wide-spread attention as a driving force in organismal evolution [1], especially in the context of contemporary debates surrounding a proposed extension of standard evolutionary theory [1–4].

Diplogastrid nematodes such as *Pristionchus pacificus* and its conspecifics are model organisms used to study the molecular mechanisms that govern phenotypic plasticity and the role it plays in phenotypic evolution [4–7]. These worms display a morphological novelty in their mouths that renders them unique amongst roundworms: they possess movable cuticular teeth which allow them to predate on other nematodes (Fig. 1A). This phenotype is plastic; depending on the environment, one of two alternative adult morphologies can be adopted: the predatory “eurystomatous” morph which has two cuticular teeth, or the strictly bacterial-feeding “stenostomatous” morph that has only a single tooth (Fig. 1A) [5,7]. Recent studies utilizing *P. pacificus* began to elucidate the complex genetics behind this polyphenism [4,8–11] and identified a modular gene regulatory network (GRN) which controls the development of plastic feeding structures found in Diplogastridae (Fig. 1B). An upstream module perceives environmental information and relays it to a central plasticity-switch module that controls which of the two alternative feeding morphologies will be developed [4,10,12]. Downstream of the plasticity-switch module is a phenotype-execution module that governs the morphogenesis of the cuticular feeding structures [4]. At the core of this module are two transcription factors, *Ppa-*NHR-1 and *Ppa-*NHR-40, which co-regulate the expression of genes involved in the modification and degradation of the cuticular material that makes up the worm’s feeding structures (Fig. 1A,B) [11]. Additionally, recent studies identified mucin-type protein DPY-6 and chitin synthase CHS-1 as members of the phenotype-execution module which participate in the synthesis of the feeding structures [13,14]. However, the transcription factors regulating their expression have not yet been identified.

**Fig. 1.**
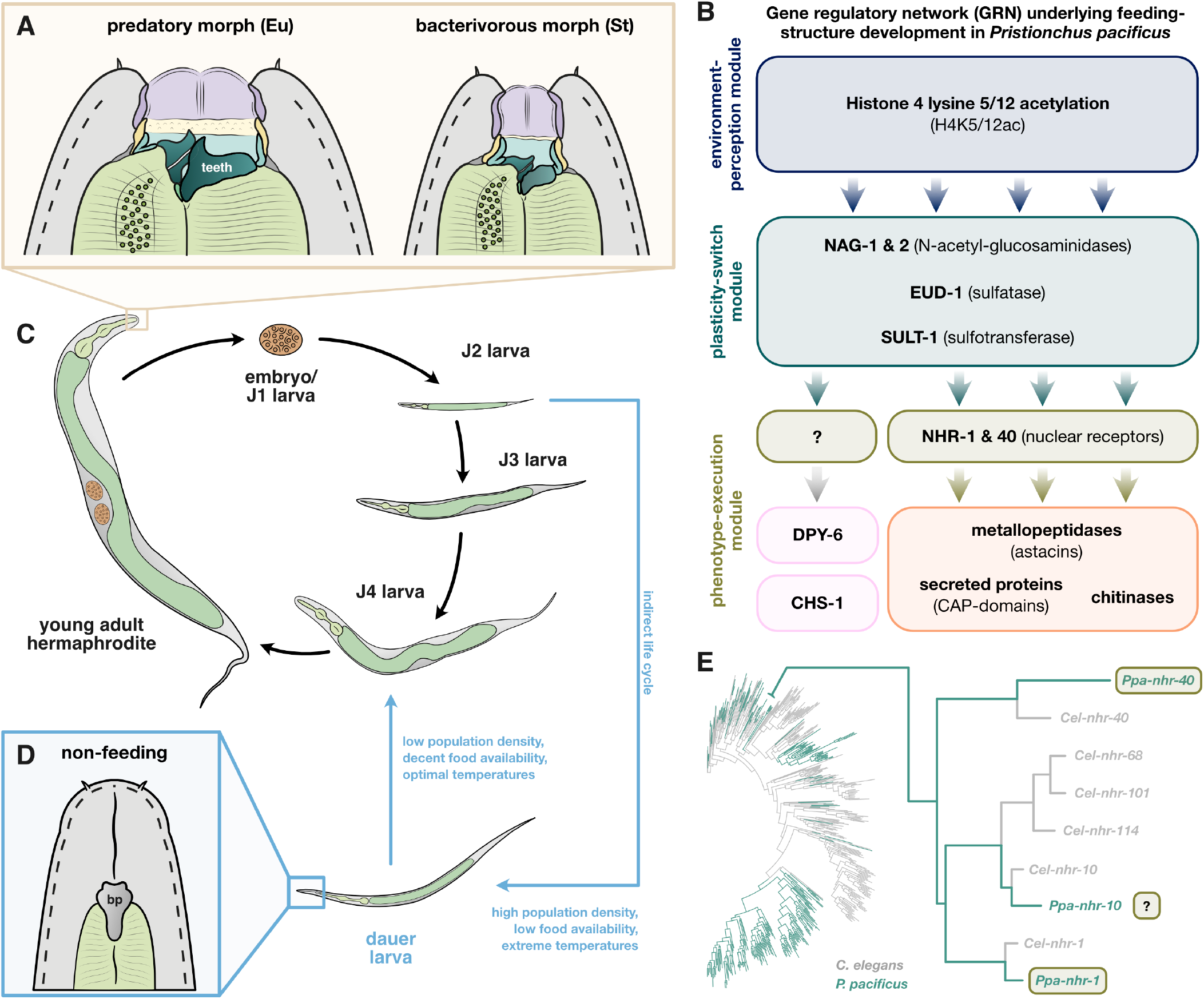
Phenotypic plasticity throughout the *Pristionchus pacificus* life cycle and the gene regulatory network of feeding-structure development. (A) Alternative adult mouth forms in *P. pacificus*. Eu = eurystomatous, St = stenostomatous. (B) *P. pacificus* life cycle, including the alternative “indirect life cycle” via dauer development. (C) Closed mouth morphology of dauer larvae. bp = buccal plug. (D) modular gene regulatory network (GRN) controlling feeding-structure development in *P. pacificus.* (E) Phylogeny of all *nhr* genes encoded in the genomes of *C. elegans* and *P. pacificus.* Tree was re-drawn based on the data of Sieriebriennikov *et al.* [11]. The asterisk (*) indicates the location of the branch that harbors the three deeply conserved sister receptors NHR-1, NHR-10, and NHR-40.

Another plastic phenotype maintained in *P. pacificus,* and other nematodes such as distantly related “rhabditid” nematode *Caenorhabditis elegans*, is related to their life cycles. Under suboptimal environmental conditions, e.g. overcrowding or food depletion, young larvae can enter an indirect life cycle (Fig. 1C) as dauers, an arrested developmental stage [15,16]. Dauers are specialized non-feeding larvae adapted for stress survival [15,17] and dispersal [18]. Characteristic morphological features of dauers include a thickened body wall cuticle and a closed buccal cavity with a plug clogging the alimentary channel (Fig. 1D) [15]. Thus, dauers cannot feed and they have to generate energy from stored neutral lipids via lipolysis and fatty acid *β*-oxidation [19–22]. Once environmental conditions turn more favorable, worms can exit dauer and resume development into reproducing adults (Fig. 1C). Interestingly, like diplogastrid feeding plasticity, various aspects of dauer plasticity are regulated by nuclear receptors (NHRs) [16,23,24], which constitute one of the most important transcription factors classes in animals [25]. They generally share a deeply conserved protein organization and contain a DNA-binding domain (DBD), which usually recognizes specific DNA motifs in the *cis*-regulatory regions of their target genes, and a ligand-binding domain (LBD), which mediates the activity state of the receptor in response to specific ligands [25,26]. Besides their roles in phenotypic plasticity [27], NHRs are known to regulate copious metabolic and physiological functions in nematodes, including immune reactions, detoxification, fatty-acids synthesis and *β*-oxidation, as well as responses to acute starvation [28–32]. Note that the majority of NHRs in invertebrates have not been deorphanized, leaving many aspects of their molecular function unknown [25–28].

Recent advances in genomics also revealed that, in contrast to other animals, nematodes often possess hundreds of *nhr* genes [11,33]. The overwhelming majority of these, called “supplementary nuclear receptors” (*supnrs*), originated from an explosive burst of duplications of the nuclear receptor HNF4 [34,35]. Another striking observation among distantly-related nematodes with similarly high numbers of NHRs (e.g. *C. elegans* [n=266] and *P. pacificus* [n=254]) showed that the vast majority of NHRs is completely lineage-specific with no one-to-one orthologs in other species (Fig. 1E) [11]. Additionally, the expression of *supnr* genes in the nervous system is highly biased toward sensory neurons, suggesting that the expansion of NHRs, which is particularly blatant in free-living nematodes, could be linked to the worm’s complex sensory environments filled with entangled microbial foods [33]. Thus, *supnrs* might act as sensors of environmental food signals or internal metabolic signals, and may adjust the worm’s physiological states or behavioral responses according to perceived environmental information [28,30,32,33].

Interestingly, *Ppa-*NHR-1 and *Ppa-*NHR-40, the aforementioned regulators of feeding-structure morphogenesis in *P. pacificus*, are representatives of these nematode-specific *supnrs* (Fig. 1E) [11] and, specifically, two of the very few *supnrs* which are actually conserved across distantly-related species. Although previous studies demonstrated that they are not only deeply conserved, but also co-express in the same pharyngeal tissues that produce the feeding structures of both *P. pacificus* and *C. elegans*, they were found to control feeding-structure development only in diplogastrids [11,36]. This can be explained by the fact that NHR-1 and NHR-40 regulate vastly different pools of regulatory target genes between these two species, and that they control very young (and rapidly evolving) cuticle-modifying enzymes only in *P. pacificus*. Together, these data suggest that these two *supnrs* were likely co-opted into the modular feeding-plasticity GRN of diplogastrids during the evolution of their tooth-like feeding structures [36].

Intriguingly, phylogenetic analyses of all *nhr* genes in *C. elegans* and *P. pacificus* found that another *supnr*, *Ppa-*NHR-10, is the sister paralog of *Ppa-*NHR-1, and that these receptors, together with *Ppa-*NHR-40, form a small cluster of deeply conserved *supnrs* in the otherwise highly species-specific canopy of the *nhr* phylogenetic tree (Fig. 1E) [11,36]. Therefore, we speculated that *Ppa-*NHR-10, too, could have been co-opted as another important regulator of feeding-structure development in *P. pacificus*, possibly even the hitherto unidentified transcription factor of those genes in the phenotype-execution module which are involved in cuticle synthesis (Fig. 1B). Thus, we set out to investigate the biological function of P*pa-*NHR-10, using an integrative approach that combines CRISPR-mediated knockouts of this receptor, geometric morphometric analyses of mutant mouth morphologies, and regulatory-target identification via transcriptomics. This study demonstrates that *Ppa-*NHR-10 is not involved in regulating mouth-form plasticity. Instead, *Ppa-*NHR-10 is required for the survival of acute starvation in young adults and long-term starvation in dauer. We also elucidated genome-wide patterns of transcriptional responses to acute starvation in *P. pacificus* and found that NHR-10’s regulatory targets include multiple enzymes involved in lipid catabolism and transport, some of which are starvation-response genes (SRGs).

## Results

### *Ppa-nhr-10* is expressed in the skin, pharyngeal glands, uteri, spermathecae, and rectal glands

Previous studies demonstrated that *Ppa-nhr-1* and *Ppa-nhr-40* are co-expressed in the pharyngeal muscle cells, the tissues that produce the cuticular teeth of *P. pacificus* [11]. Additionally, all of their joint regulatory targets (metallopeptidases, chitinases, and secreted proteins [11,36]) are expressed in a single cell: g1D. This cell is a gland cell of *P. pacificus*, which is morphologically integrated with the dorsal pharynx, where it runs through the dorsal tooth and opens into the mouth lumen [37]. We therefore wanted to determine if *Ppa-nhr-10* is also expressed in these cell types.

We generated two transcriptional reporter lines and found that *Ppa-nhr-10* is expressed in a variety of tissues (Fig. 2). These included the skin of the worms (hypodermis) (Fig. 2A, 2B, and 2C), the anterior and posterior uteri and spermathecae (Fig. 2A, 2B, and 2D), the rectal glands (Fig. 2A), and, expectedly, the pharyngeal glands (Fig. 2A and 2C). While worms consistently showed an expression in all three of the pharyngeal gland cells (g1D, g1VL, g1VR [37]), the relative intensity of the expression signal varied across the two independently acquired reporter lines: in one reporter line the ventral pharyngeal glands (g1V) dominate the signal (Fig. 2A), while in the other the dorsal pharyngeal gland (g1D) reports more strongly (Fig. 2B and 2C). Additionally, one of the lines shows only a weak expression in the rectal glands (Fig. 2A *vs.* 2B). Thus, *Ppa-nhr-10* is expressed in a variety of tissues and is not restricted to the head region.

**Fig. 2.**
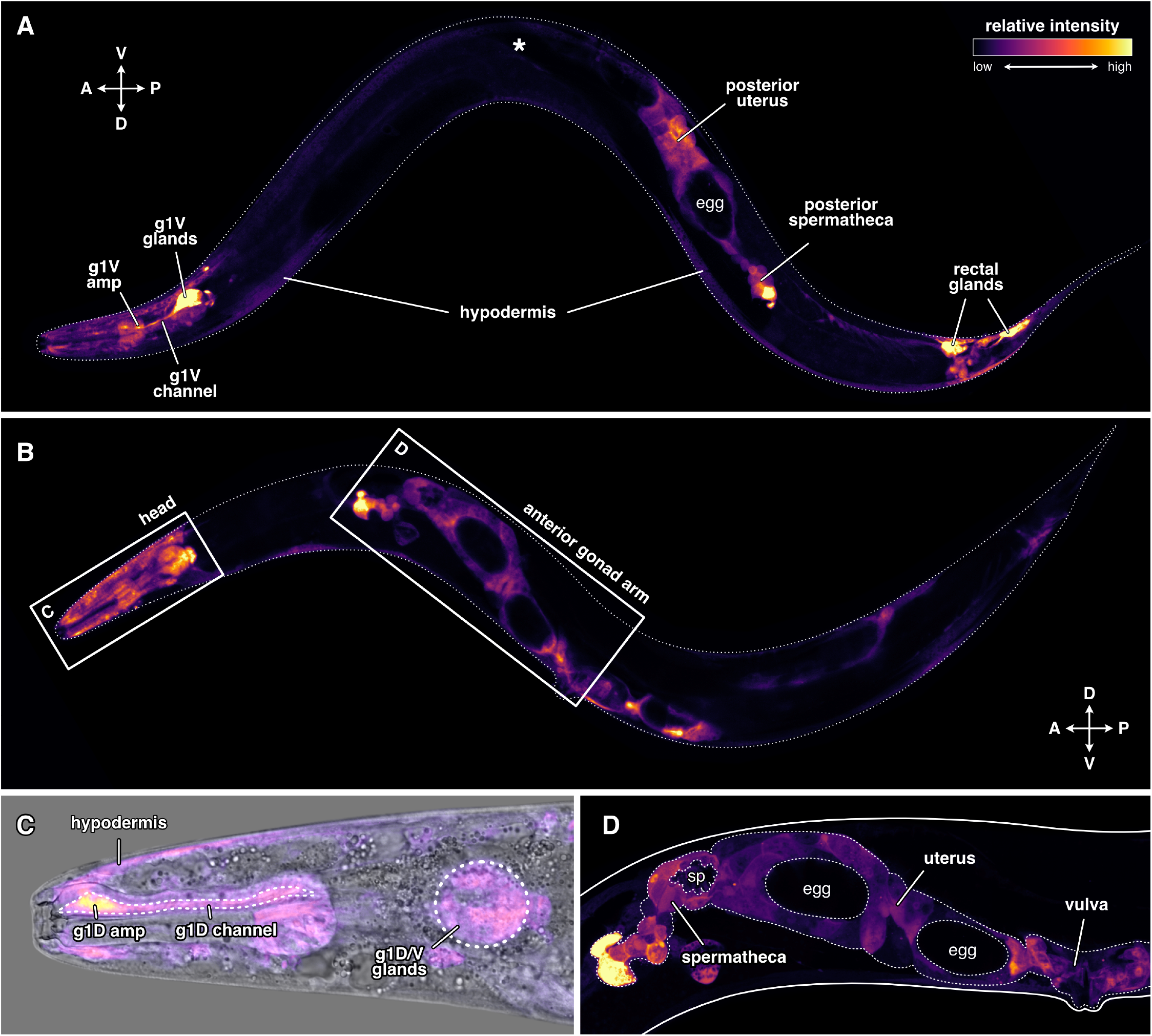
Expression pattern of *Ppa-nhr-10* across tissues. A 1.5kb long promoter sequence of *Ppa-nhr-10* (ending right before the start codon) was fused to *P. pacificus-*optimzed *GFP.* Relative strength of fluorescent-protein expression is indicated via a relative intensity scale. (A) and (B) each show a representative worm from two independent reporter lines in different orientations: (A) right body side facing reader, (B) left body side facing reader. Asterisk (*) in (A) indicates the position of anterior gonad arm. Note that both, anterior and posterior gonad arms, are always reporting; absence of the anterior arm in (A) is due to image stacking only. The crossed double-arrows with letters in (A) and (B) indicate anatomical orientation of the worm: A = anterior, P = posterior, D = dorsal, V = ventral. (C) Zoom-in on the head of the worm depicted in (B), merged with DIC image. (D) Zoom-in on the anterior gonad arm (outlined with white dots) of the worm depicted in (B). Note that neither the sperm cells nor the eggs express *Ppa-nhr-10*; only the somatic gonad tissues do so. amp = ampulla of the gland cell(s), g1D = dorsal pharyngeal gland, g1V = ventral pharyngeal glands, sp = sperm.

The fact that *Ppa-nhr-10* is the deeply conserved paralog of *Ppa-nhr-1* and *Ppa-nhr-40*, and that the expression patterns of all three receptors overlap in pharyngeal tissues associated with the morphologically novel teeth, would be consistent with the idea that this *supnr* may be another member of the GRN controlling feeding-structure development in *P. pacificus.* Thus, we set out to characterize whether the knockouts of *Ppa-nhr-10* would lead to any mouth-morphogenesis related mutant phenotypes.

### Loss of *Ppa-nhr-10* does not affect stoma morphology and only weakly affects mouth-form plasticity

Using CRISPR/CAS9, we aimed to create mutants of *Ppa-nhr-10* (genome annotation: El Paco v3, gene_ID: PPA01780, chromosome: ChrIII) by introducing small frameshift-causing insertion-deletion (indel) mutations that truncate the nuclear receptor either at the site of the DNA-binding domain (DBD) or the ligand-binding domain (LBD). While homozygous frameshift mutants in the DBD were lethal, we succeeded in generating homozygous mutant lines carrying frameshifts that truncate *Ppa-*NHR-10’s LBD (crRNA sequence: GGCGCGTGGGTTCGGGCGTA). These results indicate that DBD mutants represent lethal *loss-of-function (lof)* alleles of *Ppa-nhr-10*, and that frameshift mutations in its LBD represent *reduction-of-function (rof)* alleles that are viable. These findings are similar to previous observations in *P. pacificus* which found that certain frameshift mutations can escape non-sense mediated decay [14,38], a phenomenon that is currently under investigation in our lab. We chose two of these *reduction-of-function* mutants to establish reference alleles, *tu1654* (strain RS3920) and *tu1655* (strain RS3921), which we used for all subsequent experiments in this study (Fig 3A).

**Fig. 3.**
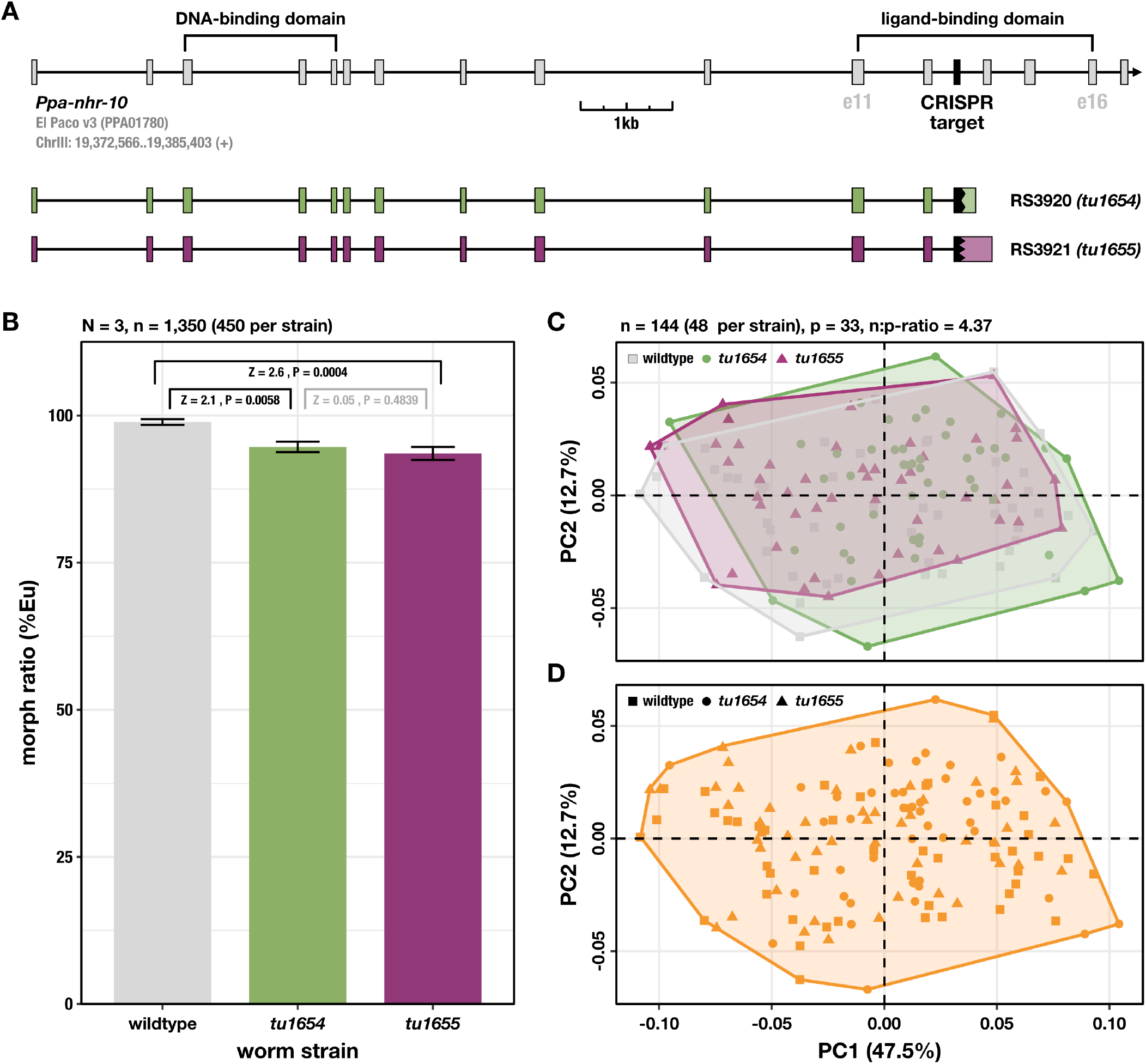
*Ppa-nhr-10* mutations do not affect mouth morphology and only weakly affects mouth-form plasticity. (A) Gene model of *Ppa-nhr-10* and frameshift alleles which lead to early truncation of the ligand-binding domain. e = exon. (B) Ratio of predatory (Eu) and bacterivorous (St) animals in the population. Worms were raised and scored on agar plates with *E. coli* OP50 as a food source. Three biological replicates were obtained (N=3) and 150 worms of each strain were scored per replicate. Significant differences are indicated in black, non-significant ones in gray. *Z*-scores and *P*-values were derived from a two-tailed PERMANOVA with null model residual randomization (RRPP). Bars represent mean values across all replicates; error bars indicate ±1 standard error of the mean. (C) Geometric morphometric analysis of form differences in the nematode mouths. The PCA plot shows the distribution of mouth morphologies of individual worms in a form space. Note that the ranges of morphological variation (outlined by convex hulls) in *Ppa-nhr-10* mutant strains completely overlap with the wild-type range of morphological variation. (D) Unsupervised (model-based) clustering reveals that all 144 individuals belong to a single morphological cluster, demonstrating that mutant worms cannot be differentiated from wild-type worms based on mouth morphology. Only predatory (Eu) animals were used for the analyses in (C) and (D).

First, we found that both *Ppa-nhr-10* mutants show a detectable decrease in the number of predatory morphs when compared to wild-type worms (Fig. 3B). While this effect is well supported by our data (wild type *vs. Ppa-nhr-10(tu1654)*: *Z* = 2.1, *P* = 0.0058; wild type *vs. Ppa-nhr-10(tu1655)*: *Z* = 2.6, *P* = 0.0004) and consistent across two alleles (*Ppa-nhr-10(tu1654) vs. Ppa-nhr-10(tu1655)*: *Z* = 0.05, *P* = 0.4839), it is only marginal in size (∼ 4-5%). This is in stark contrast to the effects observed for *loss-of-function* and *gain-of-function* mutations in *Ppa-nhr-40,* which lead to a complete loss of the predatory or bacterivorous morphs, respectively [9,11]. Given the small effect that *reduction-of-function* mutations of *Ppa-nhr-10* exert on the worm’s morph ratio, and taking into account the diversity of factors influencing the mouth-form decision [4,5,12], we conclude that *Ppa-* NHR-10 is unlikely to be a major regulator of genes involved in phenotype execution.

This idea was further supported by our geometric morphometric analysis of 2-dimensional landmark data from 144 individual worms (48 per strain), which showed homogenous and highly overlapping ranges of morphological variation in mouth form across wild-type worms and *Ppa-nhr-10* mutants (Fig. 3C). Subsequent model-based clustering of all individuals in the form dataset revealed that mutant worms could not be classified as such, or differentiated from wild-type worms, based on their mouth morphologies alone (Fig. 3D). Together, these data support the absence of any discernible mutant phenotypes in the mouth of *Ppa-nhr-10* mutants. These findings are in contrast to the phenotypes caused by frameshift mutations in *Ppa-nhr-1,* which lead to structural alterations and gives rise to intermediary mouth morphology combining anatomical features of predators and bacterivores [11].

Thus, the results of our morphological analyses do not provide support for the hypothesis that *Ppa-*NHR-10 is another constituent of the phenotype-execution module of the GRN which governs feeding-structure development in *P. pacificus*. This also indicates that not the entire NHR-1/-10/-40 cluster was co-opted into said functional context during diplogastrid evolution. Intrigued by this observation, we set out to elucidate the biological function of NHR-10 in *P. pacificus,* starting with exploratory RNA-seq experiments.

### Exploratory transcriptomics suggests *Ppa-*NHR-10 might be a regulator of lipid catabolism

Since nuclear receptors are transcription factors, it is possible to identify their putative regulatory target genes, and thus infer their potential biological functions, via comparative transcriptomics. We analyzed the transcriptomes of wild-type worms (PS312), and both *Ppa-nhr-10* frame-shift mutant alleles (*tu1654* and *tu1655*). Specifically, we performed RNAseq at two developmental stages (larvae and adults), aiming to identify potential “core targets” of *Ppa-*NHR-10 which we considered to be genes that are consistently mis-regulated in both mutant strains, in at least one of the two developmental stages.

In doing so, we were able to identify 32 potential regulatory target genes, of which 25 were down-regulated and seven up-regulated (Fig. 4). We found 11 genes consistently differentially expressed in mutant larvae and 12 in mutant adults (compared to their wild-type counterparts). The larval targets include multiple down-regulated collagens and up-regulated proteins which are known to be involved in stress responses, like C-type lectins or the UDP-glycosyltransferases (Fig. 4). Interestingly, we also found two proteins, both of them strongly down-regulated, which are predicted to be involved in the binding and transport of lipids: PPA13553 (an ortholog of the *C. elegans* vitellogenin *vit-6*) and PPA05630 (a predicted apolipoprotein). On the other hand, all of the 12 targets we identified in adult mutants were down-regulated and they include a more diverse array of proteins, ranging from a membrane-trafficking protein (PPA08133/*Ppa-scm-1*), to a component of the RNA polymerase II complex (PPA14053/*Ppa-lin-25*), to a chromatin organization modifier (ppa_stranded_DN31598_c1_g4_i2), to one of the TGF-*β* signaling receptors (PPA15180/*Ppa-daf-4*).

**Fig. 4.**
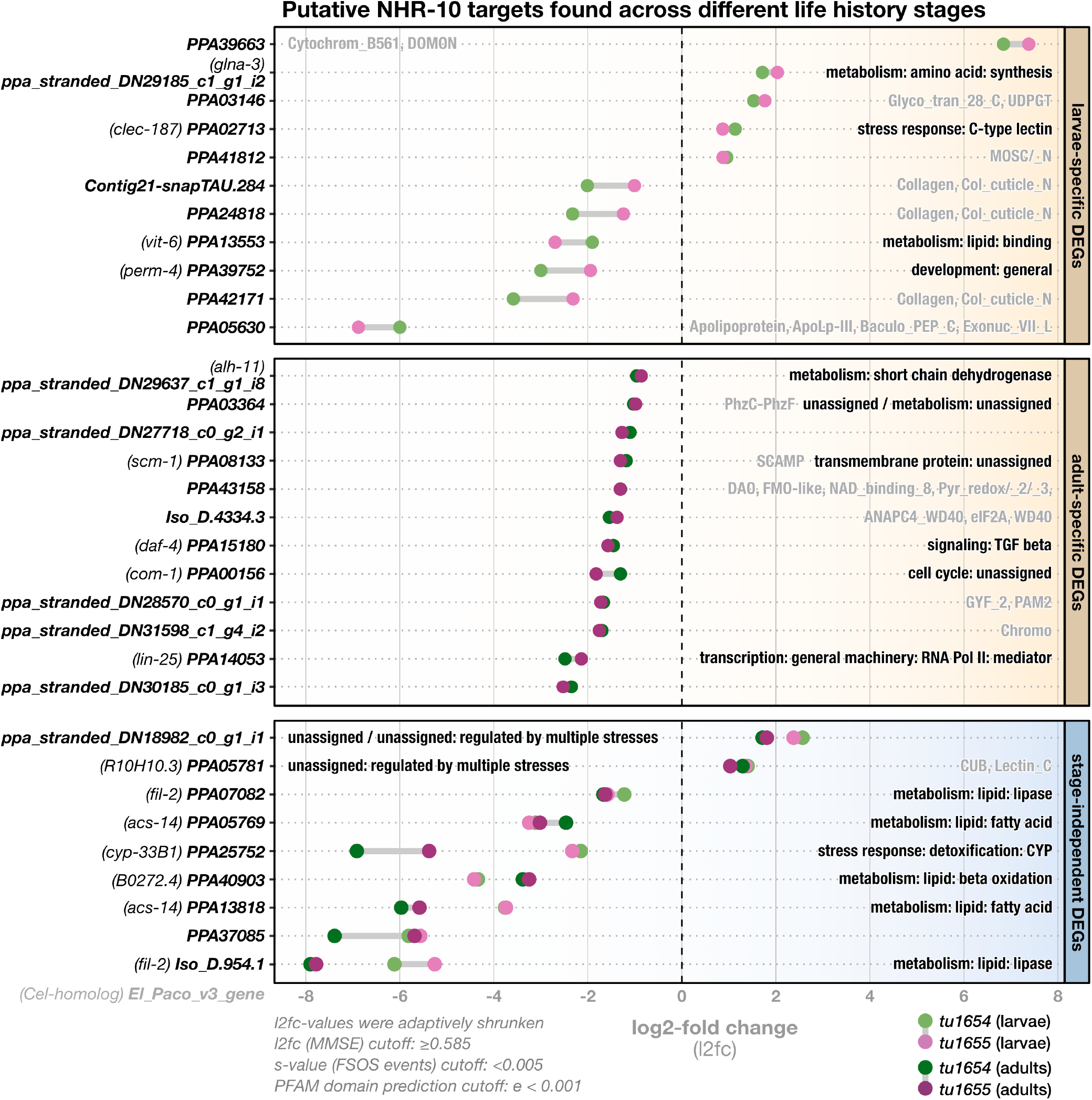
*Ppa-nhr-10* mutations affect expression levels of 32 potential regulatory target genes. Differential gene expression analysis between wild-type and *Ppa-nhr-10* mutant animals (*tu1654* and *tu1655*). Bold names along the y-axis correspond to names of the mis-regulated putative targets (El_Paco_v3). Target genes which have an exact *C. elegans* ortholog were additionally given the public gene name of this ortholog (indicated in brackets). Panels indicate in which life history stage the putative targets genes were found. Labels within the panels indicate function information for the gene in form of WormCat terms (black) and/or predicted PFAM domains (gray). WormCat annotations were adapted from *C. elegans,* based on *Cel-Ppa-*orthogroups identified with OrthoFinder. PFAM domains were only indicated for genes we could not assign a WormCat term to, or genes whose WormCat terms ended in “unassigned”. Cutoffs used in the differential gene expression analysis with DESeq2 and for PFAM domain prediction with HMMER are indicated in the lower left. DEGs = differentially expressed genes, MMSE = minimum mean squared error, FSOS = false sign or smaller.

Additionally, we found nine putative targets which are strongly and constitutively mis-regulated across all developmental stages of mutant animals, and the majority of them have predicted functions in lipid catabolism. Briefly, canonical lipid catabolism involves three processes: (I) lipolysis - the cleavage of neutral storage lipids (TAGs) into glycerol and free fatty acids by lipases, (II) activation - the ligation of CoA to free fatty acids by fatty acyl-CoA synthetases (ACSs), and (III) *β*-oxidation - the breakdown of activated fatty acids into acetyl-CoA molecules. The latter process is jointly performed by four enzyme classes: acyl-CoA dehydrogenases (ACDHs), enoyl-CoA hydratases (ECHs), 3-hydroxyacyl CoA dehydrogenases (3-HACDs), and thiolases [29–31]. Interestingly, the stage-independent putative targets of *Ppa-*NHR-10 include two predicted lipases (PPA07082 and Iso_D.954.1), two predicted ACSs (PPA05769 and PPA13813), as well as a predicted ECH (PPA40903). Furthermore, these genes are among the most strongly affected by the mutations of *Ppa-nhr-10,* which is reflected by the fact that they show some of the largest log2-fold changes (L2FC) among all putative regulatory targets (Fig. 4).

Taken together, our exploratory transcriptome and differential-expression analyses suggest that *Ppa-*NHR-10 might be a regulator of body fat in *P. pacificus.* This is evident in the observation that the expression of genes predicted to be involved in the major processes of lipid catabolism and transport is negatively affected in *Ppa-nhr-10* mutants. Given that lipid metabolism and life span are tightly linked in all animals [39], and knowing that functional interferences with any of the lipid catabolic processes in nematodes can impair their survival capabilities across various environmental conditions [29,30], we decided to quantify the survival capabilities of *Ppa-nhr-10* mutants in different conditions.

### Survival analyses reveal that *Ppa-nhr-10* is required for survival of acute starvation in young adults

In *C. elegans*, interference with the major lipid catabolic pathways can lead to altered life spans and/or rates of survival over time. While mutations in some genes of these pathways constitutively impair the worm’s survival capabilities, irrespective of environmental factors, others are known to cause such impairments only in certain conditions like starvation [29–31,39]. Therefore, we examined the survival capabilities of wild-type *P. pacificus* and *Ppa-nhr-10* mutants in two different conditions: well-fed and acutely-starved. To achieve this, we transferred young adult hermaphrodites into small cell culture flasks which either contained a standard liquid culture (LC) medium rich in OP50 or a LC medium that contained no food, and scored their survival probabilities over time until all worms in the flasks were dead.

First, we found that the obtained time-to-death data from the well-fed condition is compatible with the hypothesis that there are no differences in the overall survival capabilities of *Ppa-nhr-10* mutants and wild-type worms (Fig. 5A and 5C). Neither the global Kaplan-Meier (K-M) survival analysis of well-fed worms (Fig. 5C: χ^2^ = 5.7, *P* = 0.060431), nor the pairwise comparisons of strain-specific survival curves via a log-rank test, revealed any significant differences in survival (Fig. 5A). Yet, hazard ratios were estimated to be slightly elevated in the mutant strains (1.15 for *Ppa-nhr-10(tu1654)* and 1.13 for *Ppa-nhr-10(tu1655)*). These hazard ratios describe the risk-of-death that these mutants face at any given point of time during the experiment, relative to the wild-type level of risk. A hazard ratio of 1.0 thus indicates that mutant risks-of-death are equivalent to the wild type, and hazard ratios higher or lower than 1.0 indicate relatively higher or lower risks-of-death in mutant animals, respectively. While the hazard ratios mentioned above indicate that *Ppa-nhr-10* mutants may show a ∼15% increase in the relative risk-of-death at any given timepoint in the well-fed condition, these estimates were still found to be compatible with the null hypothesis given their associated uncertainty (Fig. 5C).

**Fig. 5.**
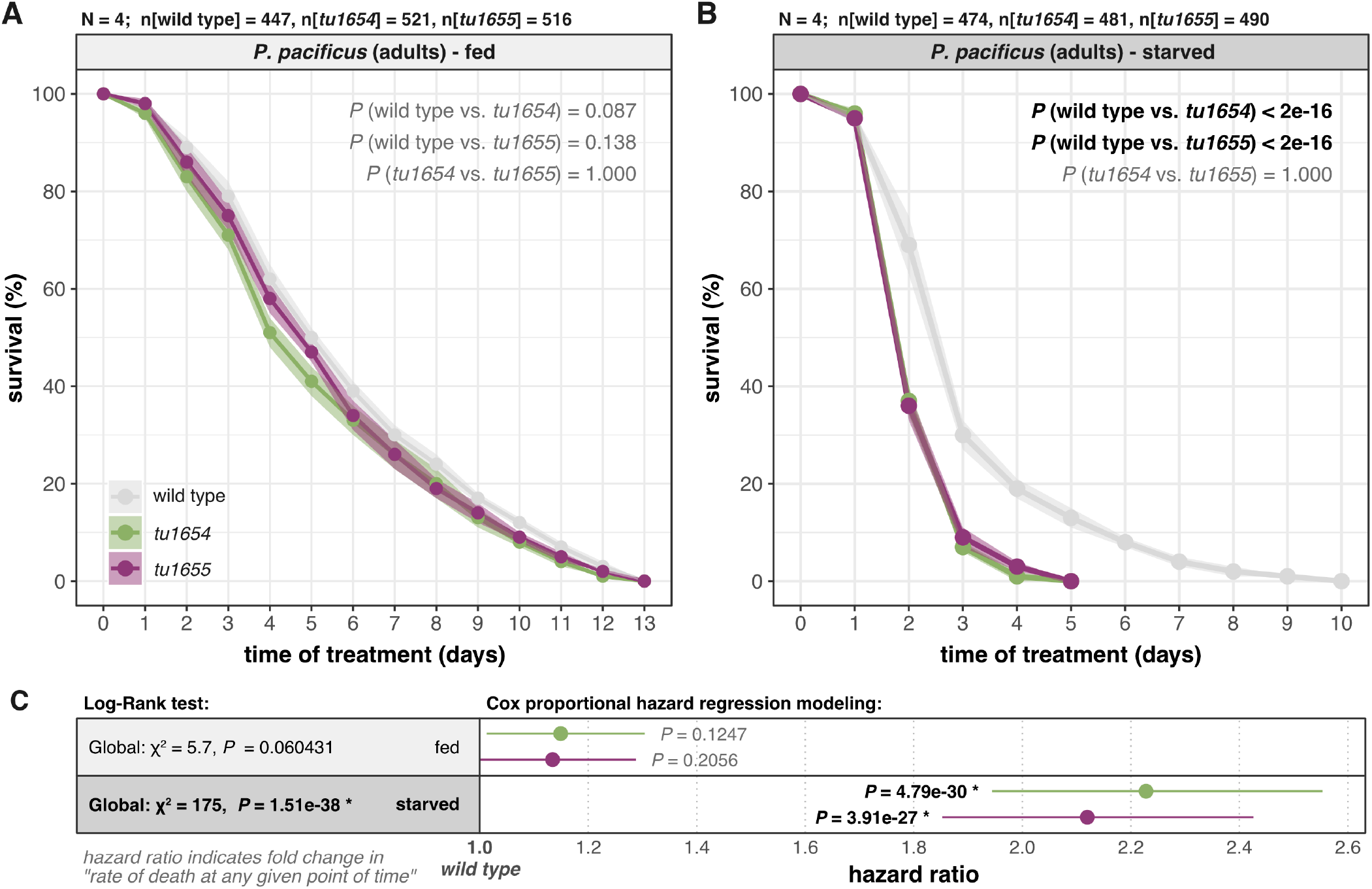
*Ppa-nhr-10* mutations cause premature death during acute starvation. Comparison of overall survival curves of wild-type *P. pacificus* and two *Ppa-nhr-10* mutants that were well-fed (A) and completely starved (B). Survival curves show the mean survival, across all replicates, over time. Transparent ribbons behind the curves indicate the associated uncertainty in form of ±1 standard errors of the mean. Each of the two experiments (A and B) was repeated four times independently (N=4); the total numbers of worms (n) that were tracked across all replicates are indicated on top of both plots. Indicated *P-*values for pairwise comparisons resulted from a log-rank test that followed a Kaplan-Meier survival probability estimation; bold and black font indicates significant differences in the pairwise comparisons of overall survival. (C) Summary of the global Kaplan-Maier statistics for both experiments (in gray boxes) next to results from Cox proportional hazard regressions (white boxes). Points indicate the estimated hazard ratios for both mutant strains within each experiment (relative to the wild-type reference level of 1.0). Intervals indicate associated uncertainty in form of 95% confidence intervals. *P*-values in bold black indicate significant differences between the mutant strain and the wild type.

Notably, we found the exact opposite to be true for the starvation experiments: the global K-M analysis indicated stark differences in the overall survival probabilities among starving worms (Fig. 5C: χ^2^ = 175, *P* = 1.51e-38) and the log-rank pairwise comparisons revealed that the survival curves of both *Ppa-nhr-10* mutants, while being virtually identical to each other (Fig. 5B: *P* = 1.0), differed markedly from the wild-type survival curve (Fig. 5B: *P* < 2e-16). Additionally, we found highly elevated hazard ratios for *Ppa-nhr-10* mutants compared to wild-type *P. pacificus* (Fig. 5C: 2.23 for *Ppa-nhr-10(tu1654)* and 2.12 for *Ppa-nhr-10(tu1655)*). These estimates indicate that for every wild-type worm that dies at any given point of time during the starvation experiment, more than two mutant worms will face the same fate. These findings strongly support decreased survival rates in mutant worms (*P*[*Ppa-nhr-10(tu1654)*]= 4.79e-30 and *P*[*Ppa-nhr-10(tu1655)*]= 3.91e-27) (Fig. 5C). While our condition-specific survival experiments revealed that acute starvation induces highly elevated rates of premature deaths in adult *Ppa-nhr-10* mutants, it is important to note that the original exploratory RNA-seq data that inspired these survival experiments were derived from well-fed worms. Thus, we wondered whether any of *Ppa-* NHR-10’s targets are starvation sensitive and therefore performed condition-specific RNA-seq experiments on well-fed and acutely-starved wild-type and mutant worms in the LC set-up.

### Starvation induces substantial transcriptional responses in genes predicted to be involved in metabolism, stress responses, and translation

To begin, we aimed to elucidate genome-wide changes of the transcriptional landscape in wild-type worms which faced 24hrs of acute starvation. Using the same differential expression pipeline as described above, we were able to identify a total of 1,007 genes showing a response to starvation (Fig. 6A). Of these, 294 genes were notably up-regulated (cluster I in Fig. 6) and 713 notably down-regulated (cluster II in Fig. 6). Thus, most of the starvation-responsive genes (SRGs) strongly decrease expression levels under starvation compared to when food is abundantly available.

**Fig. 6.**
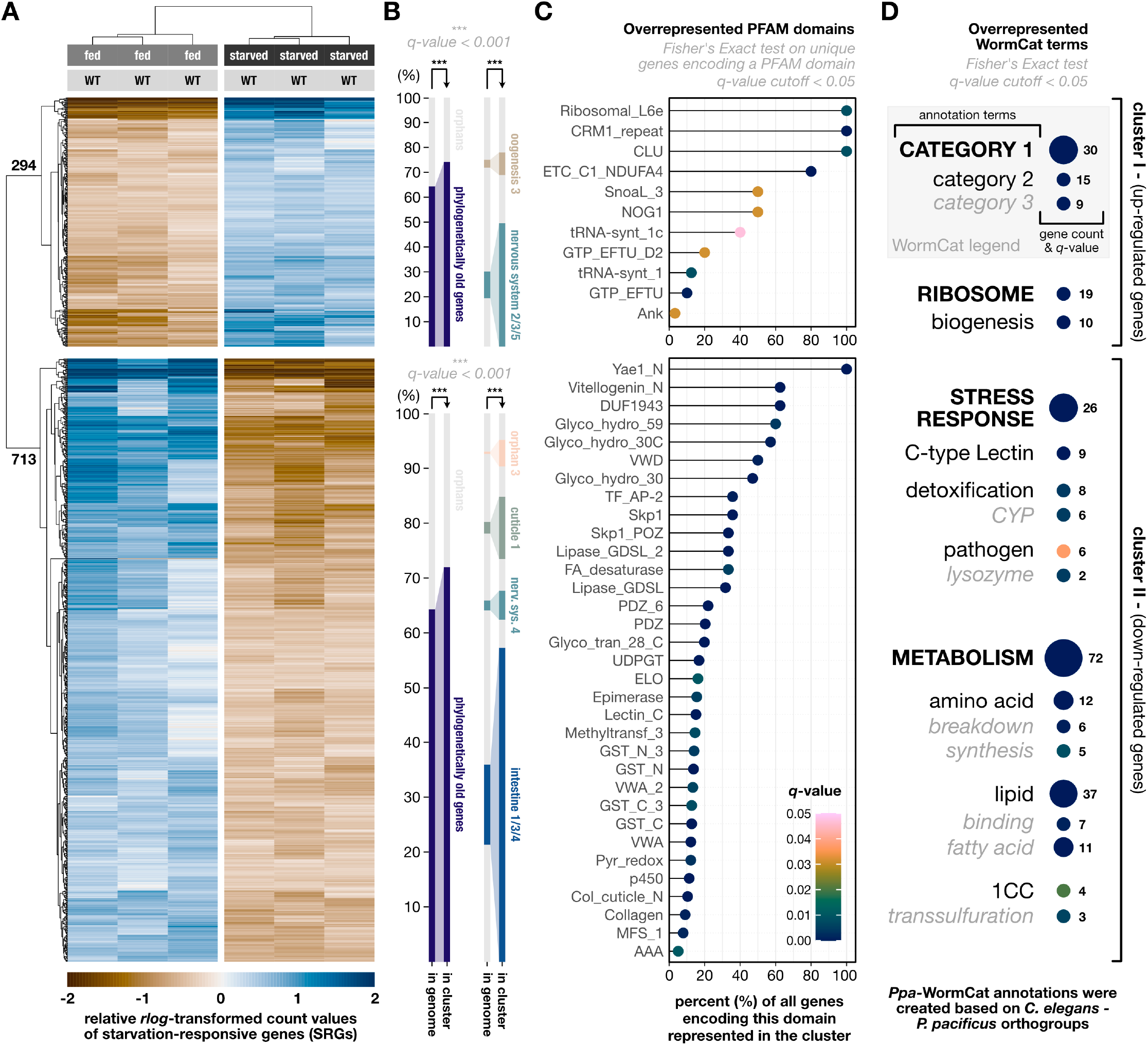
Genome-wide transcriptional responses to acute starvation in wild-type *P. pacificus*. (A) Hierarchically clustered heat map showing the 1,007 genes which were identified as differentially expressed in response to 24h of acute starvation (SRGs). Trees on the left and the top show clustering results based on similarity in expression levels. Numbers on the gene tree indicate the number of genes in each of the two response clusters. WT = wild type. (B) Bar chart showing the overrepresented gene-age classes and co-expression networks (according to [40]) identified the two response clusters, relative to the genome. “Phylogenetically old genes” refers to those genes in the genome of *P. pacificus* which are not diplogastrid-specific orphan genes. (C) Lollipop plot showing overrepresented PFAM domains in each response cluster. Y-axis gives PFAM domain names. X-axis gives the percentage of all unique genes encoding this PFAM domain, which were found to be present in the response cluster (e.g 60% means that 60% all unique genes in the genome that encode this domain are present in that cluster). (D) Bubble chart showing overrepresented WormCat terms in each of the response clusters. Enriched categories were broken down hierarchically (see WormCat legend in the upper right). Bubble size scales with the gene count indicated to the right of the bubble. The significance scale (*q-*value) depicted in (C) also applies to the bubbles depicted in (D).

We found that phylogenetically old genes, rather than diplogastrid-specific orphans, are overrepresented in both SRG clusters (Fig. 6B). Yet, while they do constitute the minority of SRGs, there are still hundreds of orphans responding to starvation (Fig. 6B). Additionally, we asked whether recently identified co-expression modules [40] were overrepresented among the SRGs. Indeed, genes belonging to various neuronal and oogenesis-related co-expression modules were strongly enriched amongst the up-regulated SRGs. On the other hand, members of intestinal and hypodermal co-expression modules were strongly enriched in the down-regulated SRG cluster (Fig. 6B).

Amongst the up-regulated SRGs, there was a clear bias towards genes encoding proteins that are critical for translational processes and ribosome assembly (Fig. 6D), including large ribosomal-subunit proteins, elongation factors, and tRNA synthetases (Fig. 6C). In contrast, genes which are down-regulated in response to acute starvation seem to control genes that predominantly encode proteins involved in diverse metabolic pathways and stress responses (Fig. 6D). The strongest detected metabolic signals originated from genes encoding proteins involved in lipid metabolism, such as fatty-acid elongases and desaturases, lipases, and vitellogenins (Fig. 6C,D). Genes involved in the synthesis and breakdown of amino acids were overrepresented among down-regulated SRGs, too (Fig. 6D). The detected stress-response signals in this cluster originated from C-type lectins, p450 cytochromes, and lysozymes (Fig. 6C and 6D).

### Condition-specific transcriptome analyses indicate that *Ppa-*NHR-10 contributes to the regulation of starvation responses

With the reference knowledge of genome-wide patterns of starvation responses as described above, we next set out to analyze the transcriptomes of starved *Ppa-nhr-10* mutants to see whether any of them are members of the identified SRGs. The condition-specific transcriptome analysis revealed an updated list of 37 putative *Ppa*-NHR-10 targets, of which 32 were down-regulated and five up-regulated (Fig. 7).

**Fig. 7.**
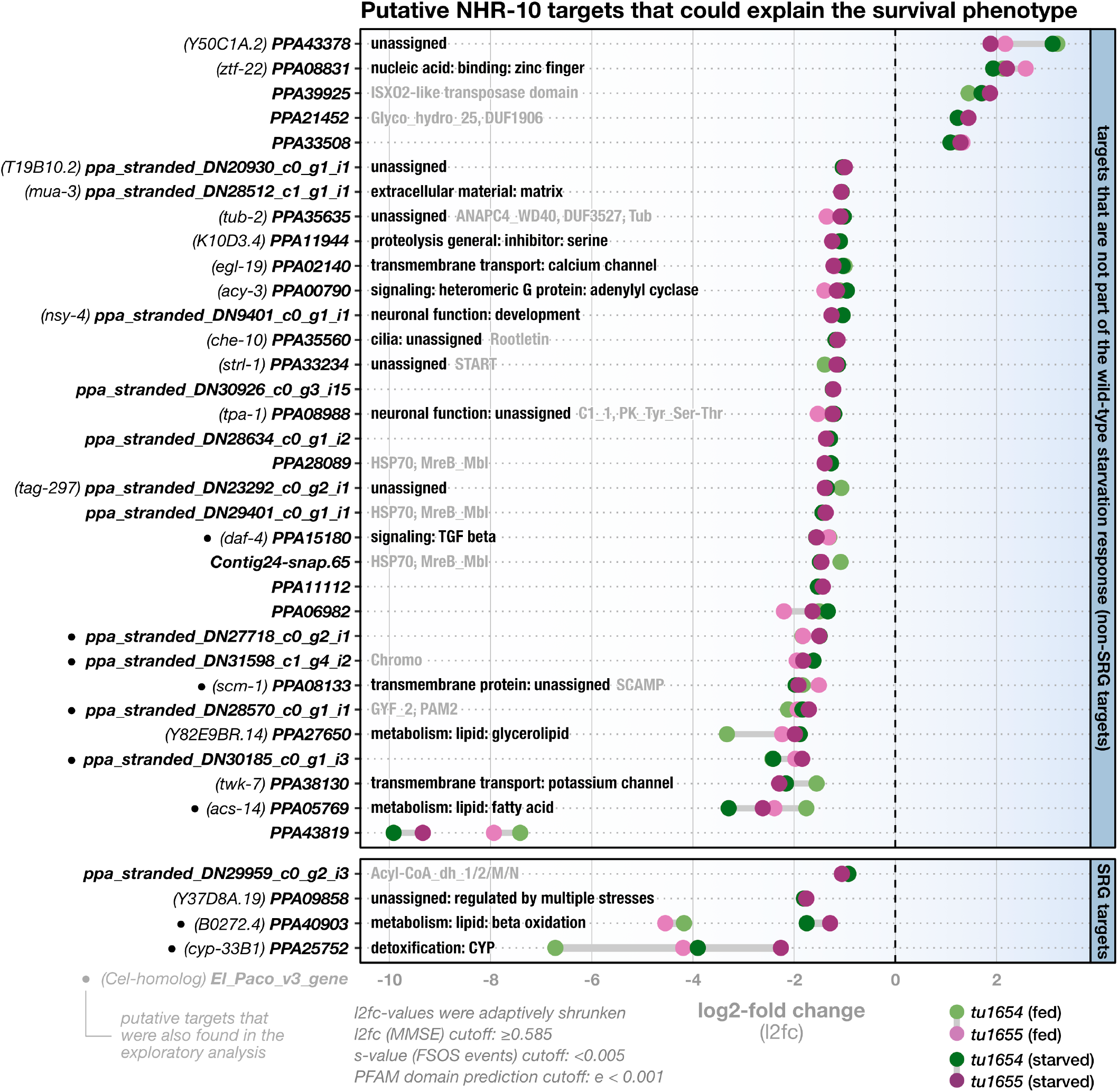
*Ppa-nhr-10* mutations disturb the expression levels of 37 putative regulatory targets in starving *P. pacificus*. Bold names along the y-axis correspond to names of the misregulated putative targets (El Paco v3) which are likely to cause the observed starvation-survival phenotype. Genes which have an exact *C. elegans* ortholog were additionally given the public gene name of this ortholog (indicated in brackets). Panels indicate whether these targets are members of the SRGs or not. Labels within the panels indicate functional information for the gene in form of WormCat terms (black) and/or predicted PFAM domains (gray). WormCat annotations were adapted from *C. elegans,* based on *Cel-Ppa-*orthogroups identified with OrthoFinder. PFAM domains were only indicated for genes we could not assign a WormCat term to, or genes whose WormCat terms ended in “unassigned”. Cutoffs used in the differential gene expression analysis with DESeq2 and for PFAM domain prediction with HMMER are indicated in the lower left. Note that, in the current El Paco v3 annotations, PPA25752 (in SRG targets) seems to be a gene model that artificially fuses two separate genes. This is supported by multiple available RNA datasets and our PFAM domain predictions, which revealed that the first half of the hypothetical protein encodes a 7TM_GPCR, while the second half encodes a p450 cytochrome. As it seems unlikely that a single protein would act as both, a GPCR and a cytochrome, we repeated our differential expression analysis after manually splitting PPA25752 into two separate genes. We found no evidence of differential expression for the resulting GPCR-gene, but could perfectly replicate the differential expression signature we originally found for PPA25752, for the cytochrome-encoding gene. FSOS = false sign or smaller, MMSE = minimum mean squared error, SRG = starvation-response gene.

The comparison of gene expression in well-fed and starved *Ppa-nhr-10* mutant animals indicates that the original exploratory RNA-seq experiments missed a large number of putative targets (28 out of 37). Many of the newly identified targets have a wide variety of predicted functions which are not easily grouped into overarching processes (Fig. 7). Yet, we recovered nine of the putative target genes that were found in the exploratory analysis (Fig. 4 and Fig. 7), including the TGF-*β* receptor PPA15180/*Ppa-daf-4,* the p450 cytochrome PPA25752/*Ppa-cyp-33B1*, and the two lipid catabolic enzymes PPA05769/*Ppa-acs-14* and PPA40903 (Fig. 4 and Fig. 7). This confirms that, while the initial RNA-seq approach missed out on starvation-specific effects, it still identified some of the targets which likely convey starvation resistance.

More importantly, the condition-specific RNA-seq approach added to the initial observation that the putative targets tend to be ones that are involved in the transport and breakdown of lipids and fatty acids (Fig. 7). PPA33234/*Ppa-strl-1* is a lipid-binding protein predicted to be mitochondrial and involved in the transport of cholesterol for steroid biosynthesis. PPA27650 is predicted to be involved in intermembrane transport of lipids as part of glycerolipid synthesis. Genes that encode important players in the *β*-oxidation of fatty acids were recovered, too: PPA05769/*Ppa-acs-14* is a fatty acyl-CoA synthetase, the type of enzyme that “activates” free fatty acids for *β*-oxidation. Additionally, the gene *ppa_stranded_DN29959_c0_g2_i3* encodes an acyl-CoA dehydrogenase (ACDH) and PPA40903 an enoyl-CoA hydratase (ECH). These two enzymes catalyze the first two of the four steps of mitochondrial *β*-oxidation [29–31].

Lastly, we wondered what the gene regulatory basis of the starvation-survival phenotype in *Ppa-nhr-10* mutants looks like and anticipated two different scenarios. One, that NHR-10’s regulatory targets would be members of genetic networks which are not part of the starvation response machinery (Fig. 6), but might, when mis-regulated, lead to decreased survival capabilities in mutant worms that face starvation. Alternatively, *Ppa-*NHR-10’s regulatory targets could actually constitute members of the SRGs. In this latter scenario, the impaired survival capabilities of starving mutants could be explained by malfunctions in the starvation-response itself. Our analyses suggested a combination of both: while most of *Ppa-*NHR-10’s putative targets do not respond to starvation, we found four SRGs among them (Fig. 7). These included the enoyl-CoA hydratase and p450 cytochrome which were already found in our exploratory RNA-seq analysis (Fig. 7). Taken together, these findings suggest that *Ppa-*NHR-10 controls some of the worm’s transcriptional responses to acute starvation.

### Dauer assays demonstrate that mutations in *Ppa-nhr-10* also affect dauer biology

Finally, we wondered whether *Ppa-*NHR-10 might play a more general role in starvation survival (beyond its requirement for starvation resistance in young adults) and investigated the effects of *Ppa-nhr-10* mutations on the development of dauer larvae (Fig. 1B). For that, we quantified three different aspects of dauer biology using a robust combination of assays: i) the number of worms which entered the dauer stage due to overcrowding, ii) the time it took the worms to exit dauer and resume development when re-introduced to food, and iii) the survival of dauers over extended periods of time (Fig. 8A).

**Fig. 8.**
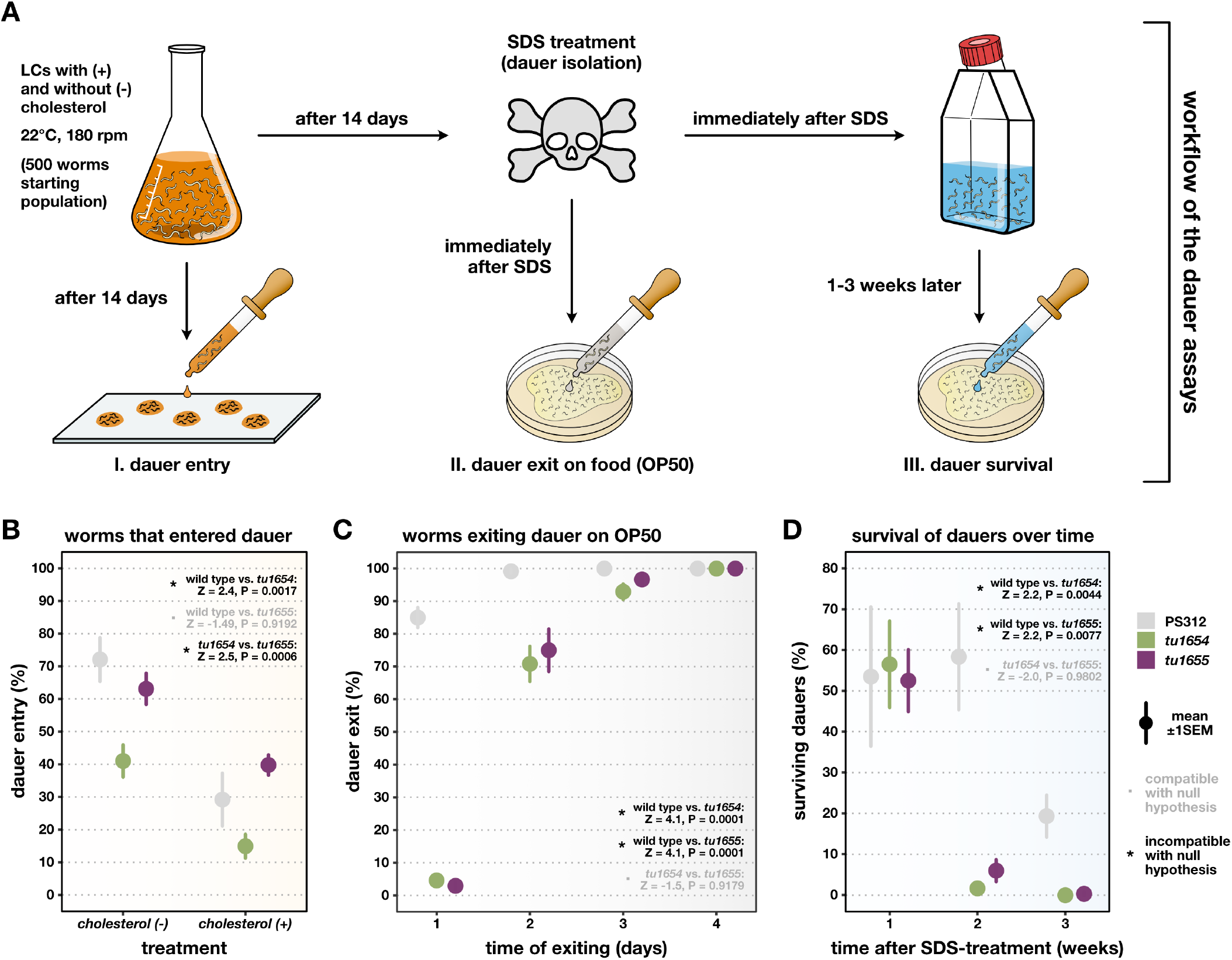
*Ppa-nhr-10* mutants have various dauer-related phenotypes, including impaired long-term survival. (A). Workflow of the three dauer assays. (B-D) Results of the individual phenotypic assays indicated above each plot in the workflow chart. Point ranges indicate the mean estimates across all replicates ±1 standard error of the mean. Note that the dauers used in the exit and survival assays (C-D) were obtained from LCs without cholesterol. Additionally, all of the worms in (C) managed to exit dauer and fully resumed developments into viable adults. Hypothesis tests used a two-tailed PERMANOVA with null model residual randomization (RRPP); relative effect sizes (*Z*-scores) and *P*-values for pairwise comparisons among the three strains are reported in the upper right of each panel. Incompatibility with the null hypothesis (*Z* ≥ 2 and *P* < 0.05) indicates the presence of a strain-specific phenotype; compatibility with the null hypothesis (*Z* < 2 or *P* ≥ 0.05) supports the absence of a strain-specific phenotype. LCs = liquid cultures, rpm = rotations per minute, SDS = sodium dodecyl sulfate, SEM = standard error of the mean. All experiments were repeated multiple times independently (N=4). Number of worms in (B): n[PS312, (-)] = 460; n[*Ppa-nhr-10(tu1654)*, (-)] = 537; n[*Ppa-nhr-10(tu1655)*, (-)] = 432; n[PS312, (+)] = 750; n[*Ppa-nhr-10(tu1654)*, (+)] = 877; n[*Ppa-nhr-10(tu1655)*, (+)] = 879. Number of worms in (C): n[PS312] = 240; n[*Ppa-nhr-10(tu1654)*] = 240; n[*Ppa-nhr-10(tu1655)*] = 240. Number of worms in (D): n[PS312] = 395; n[*Ppa-nhr-10(tu1654)*] = 175; n[*Ppa-nhr-10(tu1655)*] = 177.

Interestingly, for dauer entry, we found a mutant effect in only one of our *Ppa-nhr-10* alleles (*tu1654*). Across culture conditions, worms carrying this allele showed a significantly decreased number of dauer larvae in the overall population when compared to the wild-type animals and members of the second mutant strain (Fig. 8B). However, such allele-specific differences could not be observed in the remainder of our dauer assays (Fig. 8C and 8D). The dauer-exit data demonstrated that mutant animals consistently needed more time to exit the dauer stage and resume normal development. While all worms eventually completed the dauer exit (and developed into fully viable adults), it took *Ppa-nhr-10* mutants twice as long as wild-type worms to do so (Fig. 8C). Lastly, we found impaired long-term survival capabilities in mutant dauer larvae (Fig. 8D). For example, after spending one week in dauer, the estimated overall survival rates of wild-type and mutant dauers were indistinguishable. In contrast, after a second week, wild-type worms showed survival rates which were indistinguishable from the previous week, whereas mutant worms died off rapidly, leaving almost no survivors (Fig. 8D). This trend continued into the third week, where mutant dauers were essentially absent, while a fifth of wild-type dauers were still alive. Thus, these assays revealed that all investigated aspects of dauer biology were negatively affected in *Ppa-nhr-10* mutants. This observation confirmed the hypothesis we conceived based on the combined results from our survival analysis and our condition-specific RNA-seq data, and demonstrates that *Ppa-*NHR-10 is indeed required in the broader context of starvation resistances throughout the *P. pacificus* life cycle, including the ecologically relevant dauer stage [18,41,42].

## Discussion

In this study, we aimed to reveal whether *Ppa-*NHR-10 is a regulator of feeding-structure development in *P. pacificus* and, thus, address whether a small but complete cluster of three closely-related and deeply-conserved supplementary nuclear receptors was functionally co-opted into the gene regulatory network that governs feeding-structure plasticity in *Pristionchus* nematodes [11,36]. We were able to demonstrate that *Ppa-*NHR-10 is indeed, amongst other tissues like the skin, expressed in the relevant cell types which are part of the morphological novelties of diplogastrid mouths (e.g., the g1D pharyngeal gland cell associated with the dorsal tooth). This is intriguing, as *Ppa-*NHR-1 itself and virtually all of the regulatory targets it shares with *Ppa-*NHR-40 are co-expressed in the same cell [11]. Yet, unlike mutations in the other two receptors, frameshift mutations in *Ppa-*NHR-10 neither disrupted the worm’s ability to develop into predators, nor did it lead to aberrant mouth morphologies. These results indicate that only *Ppa-*NHR-1 and *Ppa*-NHR-40, but not the complete NHR-1/-10/-40 cluster, were co-opted when the plasticity GRN was assembled during the evolution of mouth morphological novelty in diplogastrids [36]. This also demonstrates that the genetic basis for the evolution of plastic traits and novel structures is more complicated than often presumed, and that paralogs - at least when it comes to nematode *supnrs* - might not necessarily be adequate predictors of the biological functions of genes.

Through a combination of exploratory and condition-specific mRNA-seq experiments, we were able to identify *Ppa-*NHR-10’s putative regulatory targets. These encompass a broad variety of proteins including many metabolic enzymes and multiple signaling elements (Figs. 4, 7). Interestingly, this pattern is very different from the regulatory target patterns of its paralogs *Ppa-*NHR-1 and *Ppa-*NHR-40: most of their targets belong to a single class of metallopeptidases (astacins) that is crucial for cuticle degradation and the vast majority of them are located on the X-chromosome [11,36]. This is the same chromosome that also harbors *Ppa-nhr-1* and *Ppa-nhr-40*, as well as the multi-gene locus that regulates the plastic switch decision [8,10,43]. This pattern, again, does not hold true for the loci harboring *Ppa-*NHR-10’s targets; it does not display a bias towards one particular chromosome, let alone chromosome X (Sup. Tab. 1). In contrast to *Ppa-* NHR-1’s and *Ppa-*NHR-40’s biases towards regulating classes of specific enzymes [11,36], *Ppa*-NHR-10 seems to regulate aspects of the activation and *β*-oxidation of fatty acids (Fig. 4 and Fig. 7), processes which are known to be regulated by a variety of NHRs in *C. elegans*, too [28–31]. The differences in the regulatory “behaviors” among these closely-related NHRs demonstrate why, in this particular case, *Ppa-*NHR-10’s paralogs were inadequate predictors of its function.

We also elucidated genome-wide transcriptional responses to 24hrs of acute starvation in young adult hermaphrodites of *P. pacificus*. When focusing on the broad-scale patterns in this data, three interesting observations emerge: First, during starvation, *P. pacificus* increases the expression of numerous genes which are critical elements of the translational machinery, and it decreases the expression of metabolic pathway genes, especially those affecting lipid transport, and the desaturation and elongation of fatty acids for fat storage purposes (Fig. 6C and 6D). Thus, the biological pathways affected by acute starvation are consistent between *P. pacificus* and *C. elegans* [17,44–46], indicating that the underlying starvation-response mechanisms are deeply conserved; an observation that is also reflected in the fact that most of the SRGs are phylogenetically old (Fig. 6B). Yet differences exist in the details: when *C. elegans* starves, it does not change the expression levels of ribosome-biogenesis genes, but it decreases the translation of their transcripts [46]. We speculate that *P. pacificus* might employ a slightly different strategy, in which it increases the expression of translation-related genes without increasing the translation of their transcripts. This way, the worms would not invest large amounts of energy into the protein synthesis [17], while maintaining the ability to quickly do so if food becomes available again. Future proteomic studies will be able to directly test this hypothesis and study the evolution of these molecular starvation-response strategies.

Second, the majority of up-regulated genes are associated with the nervous system and oogenesis, while the vast majority of the down-regulated genes are associated with the metabolic hub of nematodes, the intestine (Fig. 6B). Additionally, we found numerous down-regulated genes to be members of a co-expression module which is associated with the hypodermis (Fig. 6B “cuticle 1” [40]), the site of long-term lipid storage [30]. Together with the observation that lipid anabolism is down-regulated during starvation, these data could suggest that *P. pacificus*, when facing acute starvation as adults, might undergo drastic changes to their neural regulatory landscape which may, in turn, lead the worms to switch strategies from synthesizing and storing lipids to maintain their soma, to investing into egg production and ensuring population survival. This would be in line with recent predictions concerning neural regulation of body fat during starvation in *C. elegans* [30].

Third, a variety of stress responses are down-regulated in the absence of bacterial food. This includes multiple proteins predicted to be involved in the detoxification of xenobiotic compounds (p450 cytochromes), the immune response (C-type lectins), and the defense against microbial pathogens (lysozymes) [47] (Fig. 6D). This corroborates an interesting observation: the uracil-auxotrophic OP50 strain of *E. coli*, the standard laboratory food source for nematodes [48], might be slightly toxic to *P. pacificus* [49]. Upon ingesting these bacteria, the worms may have to activate some of these stress response pathways in order to be able to sustainably live off of them as a diet. Taken together, this newly-established reference knowledge of responses to acute starvation complements recent nutrigenomic advances [40] in *P. pacificus*, and it will inform future studies that may aim to evaluate the effects of different diets on the transcriptional landscape of the worm.

The present study also demonstrates that *Ppa-*NHR-10 is broadly required for starvation resistance in *P. pacificus* (Fig. 5 and Fig. 8) and that some of its regulatory targets are starvation-responsive (Fig. 7). This indicates that *Ppa-*NHR-10 is one of the probably many transcriptional regulators that contribute to the wild-type starvation response. Interestingly, the expression levels of the *Ppa-nhr-10* gene itself do not change during acute starvation (L2FC = -0.10894 and *s* = 0.79898), suggesting that *Ppa-*NHR-10 is already present in the worm’s cells before the starvation signal is sensed, and not expressed in response to starvation. Future studies may identify the ligand of this receptor and ChIPseq experiments could be used to determine its target DNA motif. This will lead to a better understanding of *Ppa*-NHR-10’s role in the starvation response, and elucidate how the food-depletion signal is detected and translated into a biological response via this *supnr*.

Lastly, we found that the loss of *Ppa-*NHR-10 negatively affected multiple aspects of dauer biology, using a unique method combining several assays. This allowed us to investigate three pertinent dauer characteristics - entry, exit, and survival - from the same initial population of dauers in conditions similar to their ecology. In its natural environment, *Pristionchus* is found primarily as dauers in soil and on its invertebrate hosts, predominantly beetles [7,18]. The ability to enter dauer is vital to ensure success in the wild, as diminishing food resources are a common occurrence. Worms that do not enter dauer under unfavorable conditions risk preserving their populations. In addition to persevering through unpredictable food availability, *Pristionchus* faces interspecific competition [41,42]. The speed of dauer exit may determine the success of a population if cohabitating with another nematode species, as first to the feast may establish, repopulate, and consume all accessible bacteria [42]. Long-term survival in dauer provides protection during extended periods of stressful conditions [18–22,24], and may play a role in evading competition while waiting for microbial populations to reestablish [42]. Therefore, starvation resistance is a trait that is crucial for the fitness of these worms in the wild: adults need to adjust to starvation and invest into the production of viable eggs, allowing future generations of worms to persist; dauers need to be able to survive over extended periods of time, allowing them to disperse and establish new populations of reproducing worms under suitable conditions.

A current limitation of our understanding of *Ppa-nhr-10* is rooted in the fact that we were unable to obtain viable homozygous *loss-of-function (lof)* alleles via frameshifts in its DBD (indicating lethality), and that the LBD-domain mutants likely represent *reduction-of-function (rof)* alleles. It is currently unclear how these two types of mutations compare in terms of their functional impact on nematode *supnrs.* Yet, frameshift mutations in the DBD and LBD of *Ppa-nhr-1* both had *loss-of-function* effects [11]. Regardless, a *reduction-of-function* could explain why we found inconsistent (that is allele-specific) effects of *Ppa-nhr-10* mutations on dauer entry. Alternatively, these could also be explained by failure to statistically detect the effect on dauer entry in *Ppa-nhr-10(tu1655)* due to a lack of power or sampling biases. This would be in line with the fact that we did not observe any other allele-specific effects in the remainder of experiments and assays throughout our study.

In summary, we demonstrated that the supplementary nuclear receptor *Ppa-*NHR-10 contributes to the transcriptional regulation of starvation responses in *P. pacificus*, and that it is required for starvation resistance in adults and dauer larvae. Based on our expression pattern analysis and transcriptome data, we speculate that this is dependent on fatty acid *β*-oxidaton in the worms’ skin - the major site of lipid metabolism [30,31]. Future molecular genetic investigations will be able to directly test this hypothesis based on the candidate targets we identified in our experiments, ultimately expanding our understanding of the physiological mechanisms that underlie starvation resistance in these nematodes.

## Conclusion

In this study, we investigated the biological function of the nuclear receptor *Ppa-*NHR-10, the sister receptor of the two known regulators of feeding-structure morphogenesis in the diplogastrid nematodes (*Ppa-*NHR-1 and *Ppa-*NHR-40). We found that *Ppa-*NHR-10 is not involved in feeding-structure development and that it regulates entirely different types of targets genes than its sister paralogs [11,36]. We showed that it contributes to the genome-wide transcriptional responses to acute starvation, and revealed that both dauer larvae (a specialized dispersal stage) and adults of *P. pacificus* are less resistant to starvation if this receptor is experimentally rendered non-functional. Thus, our findings demonstrate that these worms require *Ppa-*NHR-10 to survive some of the harsh conditions they frequently encounter in the wild [15,18–22,41,42].

## Material & methods

### Nematode husbandry

Standard protocols for the maintenance of laboratory cultures of rhabditid nematodes [48] were followed. Worms were cultured on 6cm Petri dishes with nematode growth medium (NGM). 300µl of *Escherichia coli* (OP50) were provided as a food source. Culture plates were stored at 20°C.

### Light microscopy with differential interference contrast (DIC) and morph ratios

For microscopy, all specimens were mounted on object slides with 5% Noble Agar pads which contained the sedative sodium azide (0.5% NaN_3_) and subsequently examined using a Zeiss Axio Imager.Z1 microscope with a Zeiss Plan-Apochromate 100 x 1.4 DIC objective. Image stacks were taken using a monochromatic Zeiss Axiocam 506 CCD camera. The Zen 2 Pro Software (Carl Zeiss Microscopy GmbH, 2011; version 2.0.14283.302; 64-bit) for digital microscopic analysis and image acquisition.

Relative ratios of eurystomaous (Eu) and stenostomatous (St) morphologies in each strain were scored using the aforementioned microscopy settings. Worms that had a large right ventrosublateral tooth, a hook-shaped dorsal tooth, and a promesostegostom whose anterior tips were clearly posterior to those of the gymnostom were classified as Eu morphs; animals that did not meet these combined characteristics were scored as St morphs. Statistical differences in morph ratios were assessed via a PERMANOVA (see “Dauer assays” section of the methods for details).

### CRISPR/CAS9 mutagenesis

We used CRISPR/CAS9 to create knock-out mutations of *Ppa-nhr-10*, specifically targeting their DNA- and ligand-binding domains. Standard mutagenesis protocols for model nematodes [50,51] were followed. Target-specific CRISPR RNAs (crRNAs) were generated based on the publicly available genome sequence of *P. pacificus* (www.pristionchus.org; El Paco ver. 3). CAS9 protein, universal trans-acting CRISPR RNA (tracrRNA), and target-specific crRNAs were ordered from IDT. Ribonucleoprotein complexes (RNPs) were created by combining 0.5µl of CAS9 nuclease (10µg/ml stock) with 5µl of tracrRNA (0.4µg/µl stock) and 2.8µl of crRNA (0.4µg/µl stock). This mix was incubated at 37°C for 15min and subsequently cooled down to room temperature. The RNP mix was diluted by adding enough Tris-EDTA buffer to bring the total volume of the injection mix to 20µl. In order to avoid clogging in the injection needles, the final mix was centrifuged at 14,000rpm for 2min. Injections were conducted with an Eppendorf FemtoJet microinjector at 400x magnification under an inverse Zeiss AxioVert microscope that was equipped with a Zeiss Plan-Apochromat 40x DIC objective and coupled to an Eppendorf TransferMan micromanipulator. Injections were aimed at the distal tip of the hermaphrodite gonad in young adults; animals were punctured in a 45° angle to the anterior-posterior body axis and RNP complexes were injected into the gonadal rachis. Injected P0 animals were isolated onto separate NGM plates and allowed to lay eggs for 24h at 20°C. After that, P0s were removed from the plates and F1 animals were left to grow. Upon reaching maturity, F1s were singled out to fresh plates, allowed to lay eggs for 24h and subsequently used for genotyping. DNA of candidate F1 animals was recovered via single worm lysis (SWL), target sites were PCR amplified (using the Taq PCR Master Mix produced by Qiagen), and candidate mutations were identified using Sanger sequencing (performed by Azenta Life Sciences). The F2 offspring of candidate F1-heterozygotes were genotyped in a similar fashion. Multiple homozygous mutant lines were isolated, including frameshift mutants with premature stop codons. Backups of the original homozygous mutant strains were frozen and stored at -80°C.

### Transgenesis

To visualize which cells actively express *Ppa-nhr-10*, we generated transcriptional reporter lines. We PCR-amplified a 1.371 bp long putative promoter sequence, ending right before the start codon of *Ppa-nhr-10*, and subsequently cloned it into a pUC19 vector that contained a codon-optimized GFP with *P. pacificus* introns and the *Ppa-rpl-23* 3’UTR [52]. We generally used the NEBuilder HiFi DNA Assembly Kit for Gibson assemblies, *E. coli* (NEB 5-alpha) for bacterial transformation, and InVitrogen’s PureLink HQ Mini Plasmid kit for plasmid extraction from 3ml bacterial overnight cultures. The *Ppa-nhr-10p::GFP* construct was linearized with SphI-HF (New England BioLabs) according to manufacturer’s instructions. 10 ng/µl of our linearized plasmid were injected into the gonad of the wild-type strain of *P. pacificus* (PS312), together with 60 ng/µl of SphI-linearized genomic carrier DNA and 10 ng/µl of SphI-linearized co-injection marker *Ppa-egl-20p::RFP* [52,53]. We confirmed that expression patterns were consistent across two independent reporter lines (Fig. 2A and 2B). Fluorescent worms were imaged using a Leica SP8 confocal laser-scanning microscope (cLSM).

### Exploratory RNA-seq experiments - sequencing mutant larvae and adults

To facilitate exploratory RNA-seq analysis on our *Ppa-nhr-10* mutants comparable to the RNA-seq experiments our lab previously conducted on *Ppa-nhr-1* and *Ppa-nhr-40* mutants, we adopted (with minor modifications) the sample collection and sequencing approaches that were used by Sieriebriennikov *et al.* [11]. Animals were collected for RNA extraction at 24h (mostly J2 and a few J3 larvae), 50h (J3 and J4 larvae), and 70h (young adults) after bleach-isolation of eggs. The presence and expected distribution of the desired stages at each time point was verified prior to sample collection by screening 50 specimens on each plate under a dissecting microscope. Total RNA was extracted using the Direct-Zol RNA Mini prep kit (Zymo Research) according to the instructions provided by the manufacturer. Following Sieriebriennikov *et al.*’s approach [11], we combined 500ng of RNA that was extracted at the 24h time point with 500ng of RNA extracted at the 50h time point. This sample type thus contained 1µg RNA extracted from J2-4 larvae. On the other hand, we gathered 1µg of RNA extracted at the 70h time point (adult worms) and did not mix it with RNA extracted from any other time point. Thus, we ended up with two sample types (“larval” and “adult”), which we shipped to Novogene for library preparation and mRNA sequencing. Libraries were sequenced as 150bp paired end reads on Illumina’s NovaSeq6000 platform. We sequenced the mRNA of three strains: the wild-type *P. pacificus* strain (PS312), and two independently acquired frameshift mutants of *Ppa-nhr-10* (*tu1654* and *tu1655*). Two biological replicates were collected for the larval and adult stages of each of these three strains.

### Condition-specific RNA-seq experiments - sequencing starving worms

We also performed condition-specific RNA-seq experiments, in order to identify *Ppa-* NHR-10’s regulatory targets in the same conditions in which we identified the survival phenotype of mutant animals. Additionally, we aimed to investigate the wild-type starvation response of *P. pacificus*. Therefore, using our starvation assay set-up (see “Starvation assays and survival analysis”), we sequenced mRNA extracted from young adult hermaphrodites which have been well-fed or acutely-starved for 24 hours (in liquid cultures). Just like for the exploratory RNA-seq experiments, we used the wild-type strain (PS312) of *P. pacificus* and the two reference *Ppa-nhr-10* alleles (*tu1654* and *tu1655*) for these condition-specific RNA-seq experiments. After the liquid culture treatments, we generally washed the worms three times with 0.5% Triton X-100/PBS and three times with PBS through a 20µm nylon filter net, and subsequently preformed total-RNA extractions with the Direct-Zol RNA Mini prep kit (Zymo Research), following manufacturer instructions. For each sample, ∼1µg of total RNA was extracted and shipped to Novogene for library preparation and mRNA sequencing. As before, libraries were sequenced as 150bp paired end reads on Illumina’s NovaSeq6000 platform. This time, we sequenced three biological replicates for each the three strains in each of the two conditions.

### Quantification of transcripts and differential expression analysis (DEA)

Transcript abundances were estimated from raw read files relative to the *P. pacificus* reference transcriptome using SALMON (ver. 1.5.2) [54]. Using the entire *P. pacificus* reference genome as a decoy, we built a decoy-aware transcriptome and indexed it for subsequent read mapping using an auxiliary *k*-mer hash (k=31). Reads were quantified against this index using Salmon’s *quant* command by running the program in mapping-based mode with selective alignment as mapping strategy [55]. We allowed Salmon to infer the library type, and to learn and correct for fragment-level GC-biases and sequence-specific biases. Differential expression analyses (DEA) were carried out in R using BIOCONDUCTOR (ver. 3.17) [56], TXIMPORT (ver. 1.28.0) [57], and DESEQ2 (ver. 1.40.1) [58]. Transcript-level abundance estimates generated by SALMON were summarized into gene-level count matrices using the *tximport* function. A DESeq data set was created with the *DESeqDataSetFromTximport* function and pre-filtered for transcripts which had at least ten counts across all samples. Stage-specific (exploratory analysis) and condition-specific mutation effects (condition-specific analysis) on transcript abundances were modeled with the *DESeq* function [57,58]. Obtained maximum likelihood estimates (MLE) of the log2-fold changes (L2FC) for each gene were shrunken with the *lfcShrink* function using the adaptive shrinkage estimator from the ASHR package [59]. Resulting (shrunken) minimum mean squared error (MMSE) estimates of the L2FC were reported and used to rank and visualize candidate targets. Wondering whether there is sufficient evidence in the data that a given gene is indeed at least 50% up- or down-regulated, and whether the direction of this effect was correctly estimated, we always included a L2FC-threshold of > 0.585 (absolute value) and a *s*-value threshold of < 0.005 into our hypothesis tests.

### Quantification of morphological differences in mutant animals

We quantified morphological differences in the mouths of wild-type and mutant worms, using our recently published protocol [60] which combines 2-dimensional landmark-based geometric morphometrics (GMM) with unsupervised clustering.

For this study, we obtained image stacks of the nematode mouths in lateral position and recorded the X and Y coordinates of 18 fixed landmarks which capture the overall form of the mouth using FIJI (ver. 2.1.0) [61]. All steps of the GMM analysis were performed in R (ver. 4.3.0) [62] using a combination of the GEOMORPH (ver. 4.0.5) [63–65], MORPHO (ver. 2.11) [66,67], and MCLUST (ver. 6.0.0.) packages [68,69]. We originally generated landmark data for 150 animals (50 per strain; three strains: PS312, *Ppa-nhr-10(tu1654)*, *Ppa-nhr-10(tu1655)*), performed a GPA with the *gpagen* function of GEOMORPH, and subsequently checked for Procrustes distance outliers in our original shape data. Outliers were identified separately for each of the strains of interest, using the *plotOutliers* function. Six extreme outliers were detected and removed. The outlier-corrected data set (n=144) contained landmark configurations of 48 animals per strain. GPA was performed on the outlier-corrected data to obtain a shape matrix, to which we appended the log-transformed centroid size in order to generate a form data set s*ensu* Mitteroecker *et al.* [70]. Subsequently, we ran a principal component analysis (PCA) on the form data set to visualize potential morphological differences among wild-type and mutant worms in a form space. Lastly, we assessed whether mutant worms could be classified as such (and distinguished from wild-type worms) based on their mouth morphology. For this, we used model-based clustering via the *Mclust* function. Only “meaningful” principal components (mPCs) of form variation (identified with MORPHO’s *getMeaningfulPCs* function) were used as input variables for clustering, in order to avoid overparameterization [60]. Clustering results were visualized by coloring each specimen according to cluster membership.

### Starvation assays and survival analysis

Young adult hermaphrodites (day one of adulthood) were washed off of multiple agar plates with 0.5% Triton-X100/PBS and a lazy-L scraper. The collected worm solutions were filtered through a 20 µm nylon net filter and washed three times with 1ml 0.5% Triton-X100/PBS and, subsequently, three times with 1ml PBS. The bacteria-free worm pellets were resuspended in 250µl S-medium without cholesterol and added to empty agarose plates. The S-medium was allowed to be absorbed by the agarose and the clean worms were allowed to recover for one hour. During the recovery time, liquid cultures were set up for the experiments. We used small cell culture flasks (50ml) for our assays and added a total amount of 2.5ml liquid culture (LC) to each of them. The LCs for the acute starvation assays consisted only of 2.5ml S-medium (without cholesterol). The control LC with plenty of food were set up as follows: for each 2.5ml of final LC, we centrifuged 10ml of fresh bacterial stock solution (*E. coli* OP50, OD_600_ = 0.5) for 30min at 4250rpm (4°C). The bacterial pellets were resuspended in the according amount (2.5ml per assay flask) of S-medium without liquid culture and added to the cell culture flasks. To prevent unwanted microbial contaminations, we added 1µl of nystatin and 1.2µl of streptomycin to each assay flask, irrespective of whether it contained OP50 or not. Finally, we picked the young hermaphrodites which recovered from the washing step and added them to the assay flasks (∼50 worms per flask). All assay flasks were kept in a shaking incubator set to 20°C and 130rpm and the ratio of alive worms was scored every 24hrs until all worms in the flask were dead. Media were replaced every third day in order to limit bacterial contaminations and remove offspring animals produced in the well-fed conditions. Overall differences in the survival curves within each of the two treatments were estimated using the Kaplan-Meier (KM) approach [71]. Pairwise comparisons among individual survival curves were based on the KM estimator and performed using the log-rank test [72]. Additionally, we estimated hazard rates for each of the strains in both experiments via Cox proportional hazard regression modeling [71,72]. Effect sizes were calculated as hazard ratios which compare the hazard rates of *Ppa-nhr-10* mutants (*tu1654* and *tu1655*) to the wild-type hazard rate. *P*-values were corrected for false discoveries using the Bonferroni method. All survival analyses were carried out in R (ver. 4.3.0) [62] using the SURVIVAL (ver. 3.5-5), GGSURVFIT (ver. 0.3.0), SURVMINER (ver. 0.4.9), and GTSUMMARY (ver. 1.7.1) packages [73–76]. All survival experiments were repeated four times (N=4) independently with all three strains.

### Pfam domain prediction, WormCat annotation, and overrepresentation analyses

Protein domains encoded in the proteome of *P. pacificus* were predicted using HMMER (ver. 3.3.2) [77] in conjunction with the Pfam-A.hmm database (ver. 3.1b2) [78] (hmmscan, e-value cutoff: < 0.001). The ORTHOFINDER software (default mode; ver. 2.5.4) [79] was used to define orthogroups of *P. pacificus* and *C. elegans* proteins; WormCat (ver. 2) annotations [47] for *P. pacificus* were created based on the obtained orthogroups. In a few cases, we found multiple *C. elegans* orthologs with different WormCat terms for an individual *P. pacificus* gene. In such cases, we transferred all unique WormCat terms among the orthologs to the given gene in *P. pacificus*. All overrepresentation analyses (ORAs) were based on Fisher’s Exact test; obtained *P*-values were FDR-corrected.

### Dauer assays

We performed three different assays regarding dauer biology: dauer entry, dauer exit, and dauer survival. The workflow of these assays is depicted in Fig. 8A. Starting population of 500 young adults were added to 10ml liquid cultures (LCs) with OP50 as a food source (S-medium with or without cholesterol; 4µl nystatin and 5µl Streptomycin). Worms were propagated for two weeks at 22°C and 180rpm. After that we extracted 10µl from each LC and counted the number of dauers versus the number of feeding stages in the sample to quantify dauer entry (in %). The remaining volume of the LCs was treated with SDS following standard protocols [80] to isolate dauers. Some of the obtained dauers were added to agarose plates spotted with 300µl OP50 (30 worms per plate) and followed for multiple days to quantify the speed of dauer exit. Plates were kept at 20°C and worms were scored every 24 hrs until every individual exited dauer. The rest of the SDS-isolated dauers was added to LCs for the survival assays. These LCs did not contain food nor cholesterol and were again kept at 22°C and 180rpm for up to three weeks. Every week, we extracted 75µl of worm solution and added 25µl to three OP50-spotted agarose plates. We let the worms recover on these plates for 24 hrs and then scored alive versus dead worms to obtain the ratio of survivors (in %). All experiments were repeated four times independently. For the dauer exit and dauer survival assays we exclusively used dauers isolated from LCs without cholesterol. Mutant-specific phenotypes were assessed via a PERMANOVA [81,82] with null model residual randomization (RRPP) using the *lm.rrpp* function of the RRPP package (ver. 1.3.1) [83,84]. Relative effect sizes (Z-scores) and *P-* values are reported in Fig. 8B-8D. We considered our dauer data as incompatible with the null hypothesis (i.e., “statistically significant”) if the relative effect size was at least twice as large as the standard deviation (*Z* ≥ 2.0) and the *P-*value was below a type I error rate of 5% (*P* < 0.05). If at least one of these criteria was not met, we considered the data compatible with the null hypothesis (i.e., “not statistically significant”). Null model residuals were randomized and permuted 10,000 times. Pairwise comparisons of mutant and wild-type strains were performed with the *pairwise* function of RRPP [83,84].

### Data processing, data visualization, and illustration

All data was processed, formatted, and wrangled in R (ver. 4.3.0) [62] using the TIDYVERSE [85]. All scientific plots depicted in this study were created using the GGPLOT2 package (ver. 3.4.2) [85]. Heatmaps were created using the PHEATMAP package (ver. 1.0.12) [86]. In order to make our plots accessible to people with color-vision deficiencies or color blindness [87], we exclusively used the scientifically-derived colormaps provided by the SCICO (ver. 1.4.0) [88] and VIRIDIS (ver. 0.6.3) packages [89]. Microscopic images were edited and adjusted for levels, lightness, and contrast, in Affinity Photo (1.10.5). Scientific illustrations and final figures were created in Affinity Designer (ver. 1.10.5).

## Declarations

### Ethics approval and consent to participate

Not applicable.

### Consent for publication

Not applicable.

### Availability of data and materials

All data generated or analyzed during this study are included in this published article and its supplementary information files. The raw-read data from the mRNA-seq experiments are available in the European Nucleotide Archive (ENA) repository under the accession number PRJEB65081. Upon request, all worm strains generated in this study can be provided by the Sommer lab.

### Author contributions

T.T. conceptualized the study, designed and performed most experiments, obtained and analyzed the data, created visualizations, and wrote the manuscript. T.R. and T.T. planed and conducted the dauer experiments. R.J.S. supervised the project and provided resources. All authors edited and reviewed the original draft of the manuscript.

## Acknowledgements

We thank the Light Microscopy facility at the Max Planck Institute for Biology, especially Christian Feldhaus and Aurora Panzera, for their assistance with confocal laser-scanning microscopy. We are grateful to Heike Haussmann for freezing samples of all the worm strains produced for this study. Additionally, we thank Christian Rödelsperger, Adrian Streit, and Devansh R. Sharma for helpful discussions of our work.

---

**Sup. Tab. 1.** Overrepresentation analysis shows no bias towards any given chromosomal location among *Ppa-*NHR-10’s targets. The 37 candidate regulatory targets (DEGs) of *Ppa-* NHR-10, which were identified in the condition-specific RNA-seq experiments, were used as input for the analysis. No significant bias (FDR-corrected *P*-value < 0.05) towards any particular chromosome was found for *Ppa-*NHR-10’s target genes. DEG = differentially expressed gene, FDR = false discovery rate.

| chromosome | number_of_genes | number_of_DEGs | fold_enrichment | FDR_corrected_pvalue |
| --- | --- | --- | --- | --- |
| Chr1 | 6869 | 5 | 0.5684764688986556 | 0.35319808452624085 |
| Chr2 | 4462 | 11 | 1.9253031606236448 | 0.13589186241137188 |
| Chr3 | 3736 | 5 | 1.045199374963829 | 0.9702253905888654 |
| Chr4 | 6594 | 10 | 1.1843690824582545 | 0.8351614949834785 |
| Chr5 | 4097 | 5 | 0.9531034573748757 | 1 |
| ChrX | 3002 | 1 | 0.26015089039739275 | 0.35319808452624085 |

